# Machine learning can identify orphans that have diverged into the “twilight zone” of sequence similarity

**DOI:** 10.1101/2025.06.06.658213

**Authors:** Emilios Tassios, Jori de Leuw, Christoforos Nikolaou, Anne Kupczok, Nikolaos Vakirlis

**Author notes:** These authors contributed equally (Emilios Tassios, Jori de Leuw).

## Abstract

Species-specific orphan genes (orphans) lack homologues outside of a given taxon and frequently underlie unique species traits. It is thus important to elucidate the evolutionary origins of orphans. Orphan genes can result from sequence divergence beyond recognition, when homologous proteins diverge to an extent at which tools that rely on sequence similarity to establish homology can no longer identify them as homologues. Orphans can also result from other processes, including de novo gene emergence from previously noncoding sequences, in which a homologous protein-coding gene truly does not exist.

Here we propose that orphans resulting from divergence might be recognizable from their patterns of non-statistically significant similarity hits which are almost always discarded. To test this, we simulated diverged orphan protein sequences based on conserved proteins from the Unified Human Gastrointestinal Protein catalogue (UHGP) and used reversed protein sequences as negative data sets. We trained four machine learning classifiers on features extracted from the similarity search tool DIAMOND’s output, like total query coverage or maximum bit score. We tested the influence of evolutionary parameters such as simulation tree branch length, indel rate and among-site rate heterogeneity.

We found that the performance of the models depends on the simulation parameters: when the underlying simulated divergence was moderate, accuracy reached ∼90%, but when extremely diverged scenarios where simulated accuracy dropped to ∼70%. The most important features for the classification were the number of alignments (hits) and the minimum hit E-value. When applying our classifier on a set of ∼170,000 eligible simulated orphans from the UHGP dataset, we found that ∼30% of them are predicted to be divergent and these are shorter and more disordered than the rest. Our classifiers and pre-processing python scripts are available at https://github.com/emiliostassios/Classification-of-divergent-genes-using-ML and can be readily used as a computationally fast means to obtain a candidate set of diverged orphans from any similarity search output. Therefore, our work allows to study such orphans across the tree of life and in doing so to recover cases of remote homology, get better estimates of the evolutionary age of protein families, detect cases of rapid divergence and more generally better understand how genetic novelty arises.

## Introduction

When the first eukaryotic chromosome, chromosome III of the budding yeast *S. cerevisiae*, was fully sequenced, one of the most striking findings was that half of the protein coding genes had no detectable homologues in other lineages^1^. As more and more genomes were sequenced and became publicly available, this proportion was reduced yet it became clear that genes with seemingly no shared ancestry exist in both eukaryotes and prokaryotes, accounting for even up to one third of genes in a genome^1,2^. Orphans, as these were called, were and remain intriguing not only because their evolutionary origins are puzzling, since they do not seem to be the result of reuse and rearrangement of existing genetic components^3,4^, but also because of several distinguishing properties that they share, including fewer exons and overall shorter length than conserved genes^2,5^.

Orphans matter because they frequently underlie unique phenotypic traits^6,7^. The octopus has reflectin genes that make them capable of rapidly becoming invisible to enemies and prey^8^. The water-strider insect has unique genes that have changed the morphology of its legs to allow it to live and hunt at the surface of fast-flowing water^9^. In arctic gadid fish, an antifreeze glycoprotein has evolved de novo and provided a life-saving trait allowing to survive and adapt to subfreezing temperature water^10,11^. And the ocean is filled with an “ocean” of novelty: thousands of uncharacterized genes specific to that habitat, discovered in metagenomic samples^12^. It is clear that genomic novelty can produce phenotypic novelty and drive adaptation to new environments^6,13^. Some examples are especially intriguing: evidence suggests that the evolution of eusociality in insects, a major transition in the history of life, went together with a massive arrival of novel genes^14^. Many primate-and human-specific genes have been found upregulated in the brain^15–17^, suggesting a possible role in the evolution of important human-specific traits.

When a gene is found to be without homologues outside a particular lineage, assuming sufficient genomic data on outgroups are available, there are two main possible explanations^18^. The first is that the gene has homologues but their sequences have diverged beyond recognition, into the “twilight zone” of sequence similarity. When this happens, similarity search tools like BLAST can no longer detect them ^2,18^. This can occur when genes are under weak purifying selection, for example because of redundancy following a duplication event^19^ or due to environmental change, when genes undergo adaptive accelerated evolution^20^, or, if the encoded protein structure is flexible enough. Though this scenario seemed as the most probable (and should be the null hypothesis), studies have estimated that it may not explain the majority of genes without similarity. In the past we have used conserved microsynteny to show that, on average, approximately a third of orphan genes in three eukaryotic genomes can be attributed to divergence^18^, while others have used a simple mathematical model to estimate this percentage as slightly higher^21^. The second possibility is that the gene has evolved de novo, in which case there are truly no homologous protein-coding genes to be found. De novo gene birth is the process by which a gene emerges from a previously non-coding region of the genome^22,23^. These genes are entirely novel and by definition share no homology with any other gene^24^. Note that the above scenarios can also apply only to a part of a gene (e.g. by an existing gene extending its CDS into its 3’ region) and can be combined with other events such as gene loss in a specific lineage, domain recombination, gene fusion etc.

Can these two cases of gene origination be distinguished? In other words, is it possible to “reunite” a pair of homologues that has diverged into the “twilight zone” of sequence similarity? Once BLAST searches fail to detect any statistically significant similarity, Hidden Markov Model (HMM) profile-based searches constitute one option for remote homology sensing^25^. Yet for orphan genes that are restricted to only a single species, as is often the case, the power of HMM profile searches is limited by the fact that we cannot build inter-specific alignments for them. Thus far, several studies have studied the evolutionary origins of orphan genes in various organisms, from plants, to flies and primates^26–34^, as well as some in viruses and prokaryotes^35–37^. Most of them were based on comparative genomics using sequence similarity algorithms with various cut offs and synteny-based approaches, and a few works also employed HMM profile-based searches^29,38^. However, with the exception of the two studies mentioned in the previous paragraph (employing conserved microsynteny^18^ and mathematical modelling^21^, and ^39^ which relied on the latter), we are not aware of other works proposing new methodologies to identify diverged orphans. It is important to note that the structure prediction revolution driven by AlphaFold^40^ and the structure-based homology searches it has enabled^41^ are a substantial advance, yet the lack of alignments limits its use for species-specific genes and, in general, both alignment and single sequence prediction tools seem to perform poorly on such proteins^42^.

While orphan genes have, by definition, no statistically significant similarity hits outside a given taxonomic group, they often have non-statistically significant ones. We hypothesized that, for orphans that are the result of sequence divergence, such hits might harbor distinctive patterns that can be discriminated from random noise. We thus set out to test this novel idea using classical machine learning models. Motivated by the general paucity of studies on orphan genes in prokaryotes, we took advantage of our recent work that identified more than six hundred thousand species-specific orphans within the Unified Human Gastrointestinal Protein (UHGP) catalogue which represents the entire diversity of human gut bacteria^35^. We simulated the evolution of divergent orphans using conserved proteins and trained four classification models with features extracted from DIAMOND^43^ searches, to identify the simulated ‘orphans’ from the negative controls which were the reversed counterparts of the simulated sequences.

## Results

### Overview of the simulation-based machine learning approach

To classify orphan genes based on their similarity search output, our first step was to generate a large number of orphan sequences via simulated divergence. We developed a custom pipeline to perform this task (**Figure 1**): first, we randomly sampled the UHGP50 conserved protein clusters to obtain subsets of equal size. These were used as input for the sequence evolution algorithm with different parameters for each one of them (see **Methods**). Simulations were conducted with phyloSim^44^, which takes as input a set of sequences, a set of parameters including a substitution matrix, and a phylogenetic tree along which the simulation takes place. The simulated sequences were aligned against the UHGP50 with DIAMOND^43^, and those that had no statistically significant hits (i.e., no hits with e<0.001) were characterized as simulated orphans (referred as such in the rest of the manuscript).

**Figure 1.**
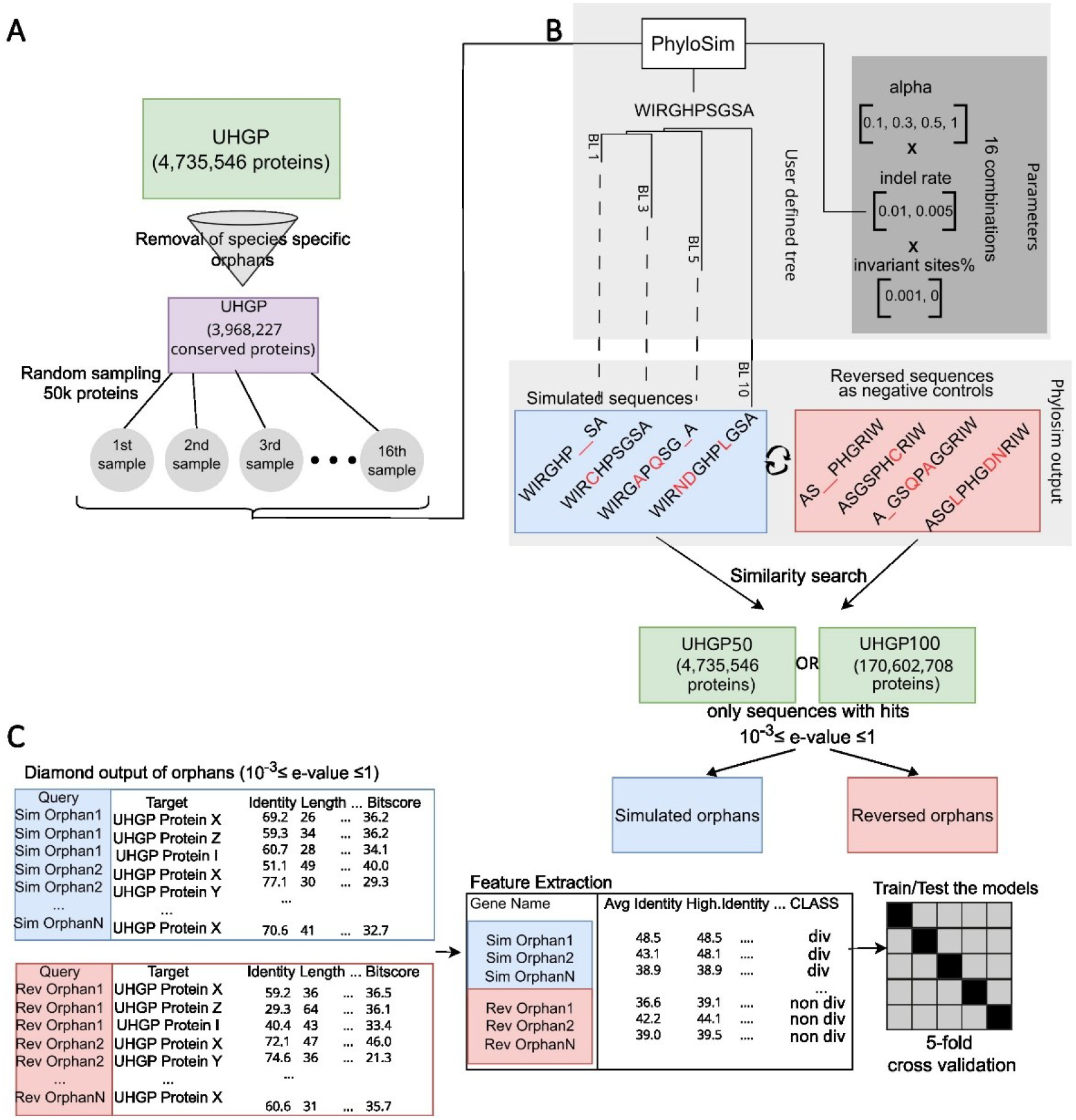
Overview of the simulation-based pipeline. **A**) We used only the conserved sequences of the UHGP database to create 16 different subsets of 50,000 thousand proteins that were used as template sequences for sequence evolution simulations. Different parameter values were assigned to each subset. **B**) The R package phyloSim was used to conduct the sequence evolution simulations. The input of the tool was i) a user defined tree, ii) a substitution model (LG), iii) a set of sequences, iv) values for parameters (alpha value, indel rate, proportion of invariant sites). We created negative controls set for each subject of simulated orphans by reversing their sequences. DIAMOND was used for similarity search of both the simulated orphans and the negative controls back to the UHGP50 or UHGP100 database. **C**) We extracted the DIAMOND features from the similarity search output and used them as input to train and test 4 classification models.

We generated 16 different sets of 50,000 sequences each that cover a range of simulation parameters, namely alpha value for rate distribution across sites, indel rate and proportion of invariant sites. We used a tree with 4 terminal branches, each with lengths of 1, 3, 5, and 10, along which the simulation was conducted.; phyloSim outputs as many simulated sequences as the terminal branches of the tree provided (in our case that is 4), thus we obtained 3,200,000 simulated sequences in total. For a simulated orphan to be included in downstream analyses, it needed to have at least one non-significant DIAMOND hit with 10^−3^ ≤ E-value ≤ 1 in the similarity search against the UHGP50 database.

As a negative set, we used reversed simulated sequences as this is known to result in sequences with the same composition but no true homologs in the database: as proteins do not evolve by reversal, any similarity will be the result of chance^45^. We implemented the exact same pipeline on the reversed set as we did with the simulated sequences, meaning that we used only the ones that did not have a significant similarity hit with any sequence from the UHGP50. The similarity search output of both simulated orphans and reversed ones were used to extract simple statistical features with which we trained a Bayesian, a Logistic Regression, a Gradient Boosting and a Random Forest classifier (see **Methods** for full details). For each sequence, we generated the following features based on hits with 10^−3^ ≤ E-value ≤ 1: number of alignments (hits), minimum E-value, average identity, average E-value, average bit score, highest percentage identity, average alignment length, highest bit score, alignment length and total query coverage.

UHGP50 contains one representative protein for each family clustered at 50% identity by Almeida et al, which means that similarity searches using simulated sequences can basically only be expected to recover low similarity homologues apart from the root sequence itself (since sequences with >50% similarity are absent). To understand whether our approach works when including all available homologues, we also generated a dataset of identical size using the UHGP100 database as a target for similarity searches, in which only 100% identical sequences are collapsed into clusters (database size=170,602,708 sequences). We thus test our approach in two scenarios: one without any close homologues (UHGP50), and one with close homologues (UHGP100). Hereafter, we present results from UHGP50-based similarity searches in our main figures with the equivalent for UHGP100 for key findings in supplementary figures.

### Strong classification performance for scenarios of low to moderate simulated divergence

In our simulation, orphan sequences result from the accumulation of mutations which in turn are governed by the simulation parameters. We observed substantial differences in the numbers and percentages of eligible orphans (i.e. with only non-significant matches) between different sets of parameters (**Figure 2A, Supp. Figure 1A**), where alpha had the largest effect. The sets which were simulated under alpha values of 1 had a total of 131,838 orphans out of 800,000 root sequences (16.4%). In contrast, simulations with alpha = 0.1 produced ten times less orphans, 13,142 (1.6%) with the same number of root sequences. Overall, there is a correlation between alpha and number of orphans (R^2^=0.27, p-value=1.248e^-5^) (**Supp. Figure 1B**). Under lower alpha values, the gamma distribution of substitution rates among sites tends to be L-shaped, leading to most sites having low rates^46^. Such sites can then form segments where similarity persists and can be detected by DIAMOND which in turn leads to fewer orphans compared to higher alpha values, where most sites have moderate rates. Indel rate had a smaller albeit significant overall effect on the number of eligible orphans, as a value of 0.01 resulted in 99,365 orphans, whereas a value of 0.005 resulted in 196,219 orphans (**Supp. Figure 1A**). We observed no effect for the percentage of invariant sites (**Supp. Figure 2**). The number of orphans varied strongly with branch length used across all datasets (**Figure 2A;** same results when using UHGP100, **see Supp. Figure 3A**). The shortest branch had a length of 1 and for all the sets combined 3,602 orphans were produced whereas from the longest branch, length 10, a much larger number of orphans was produced (n=155,652). Note that this difference in numbers is due to more sequences without any statistically significant hit in the longer branches (i.e. more sequences without any hit with E-value ≤ 0.001), and not due to differences in number of sequences with at least one non-statistically significant hit (1 ≥ E-value ≥ 0.001). Another difference due to the simulation branch lengths is that orphans simulated on the shorter branches were shorter than those simulated on the longer ones. (**Supp. Figure 4**).

**Figure 2.**
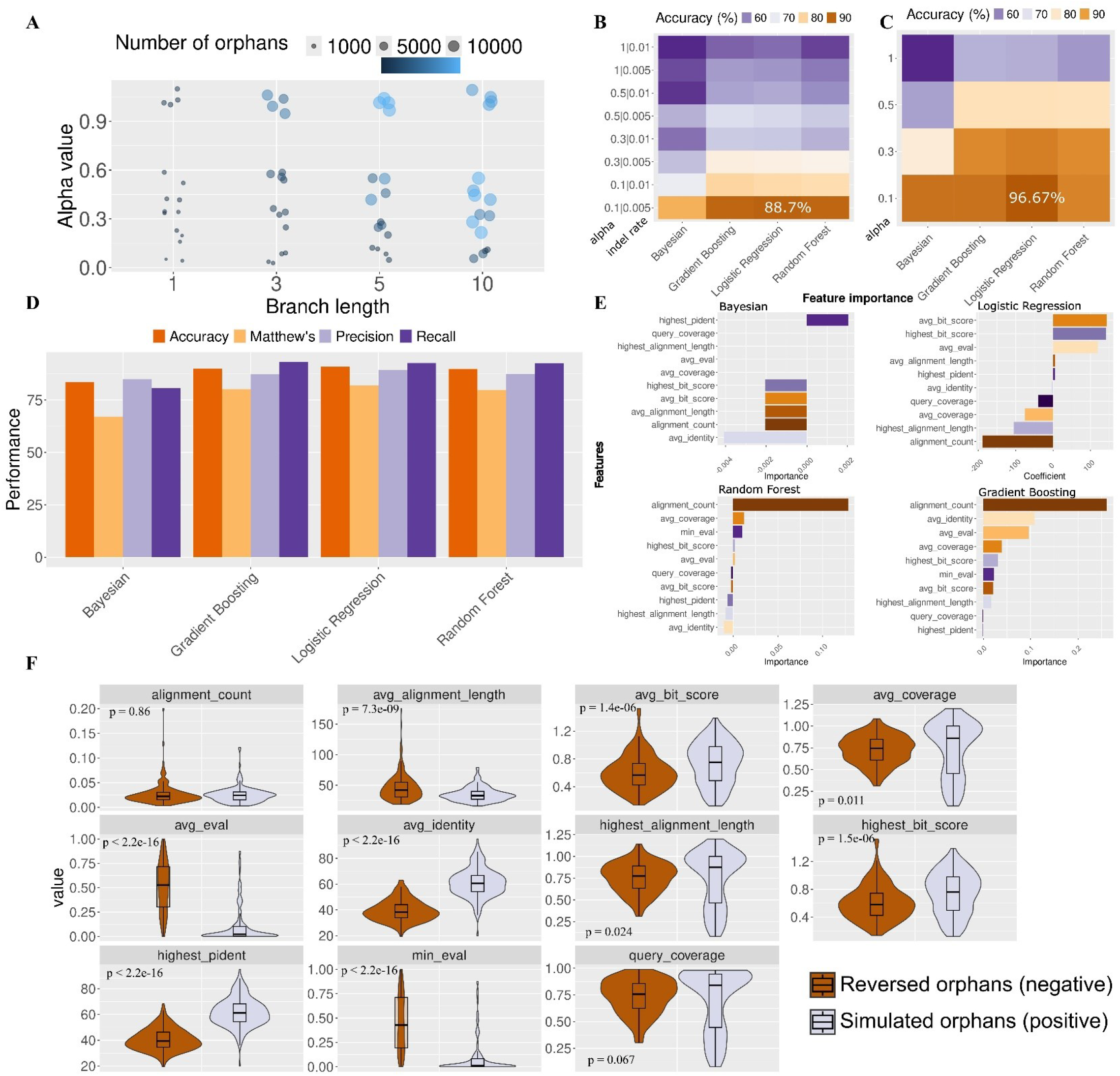
Classification performance at predicting diverged orphan sequences. **A**) Number of simulated orphans per alpha value and branch length. **B**) Accuracy metric for all four models (Bayesian, Logistic regression, Gradient Boosting and Random Forest) for all subsets produced with various combinations of alpha and indel rate values, without invariant sites. Versions with invariant sites (value=0.1) are not shown here for simplicity, as the effect of this parameter negligible. **C**) Accuracy metric for all four models and alpha values, using sequences that were simulated without indels. **D**) All performance metrics (accuracy, recall, precision, Matthew’s correlation coefficient) for the best performing models (simulated with: alpha = 0.1, indel = 0.005, inv_sites = 0.001; only simulated orphans from the two shortest branches used), after correcting for length differences between simulated and reversed orphans. **E**) The most important features for each of the best performing models. **F)** Feature comparison between the reversed orphans (negative) and simulated divergent orphans (positive) for the dataset used in the best performing models. Wilcoxon test P-values are shown.

**Figure 3.**
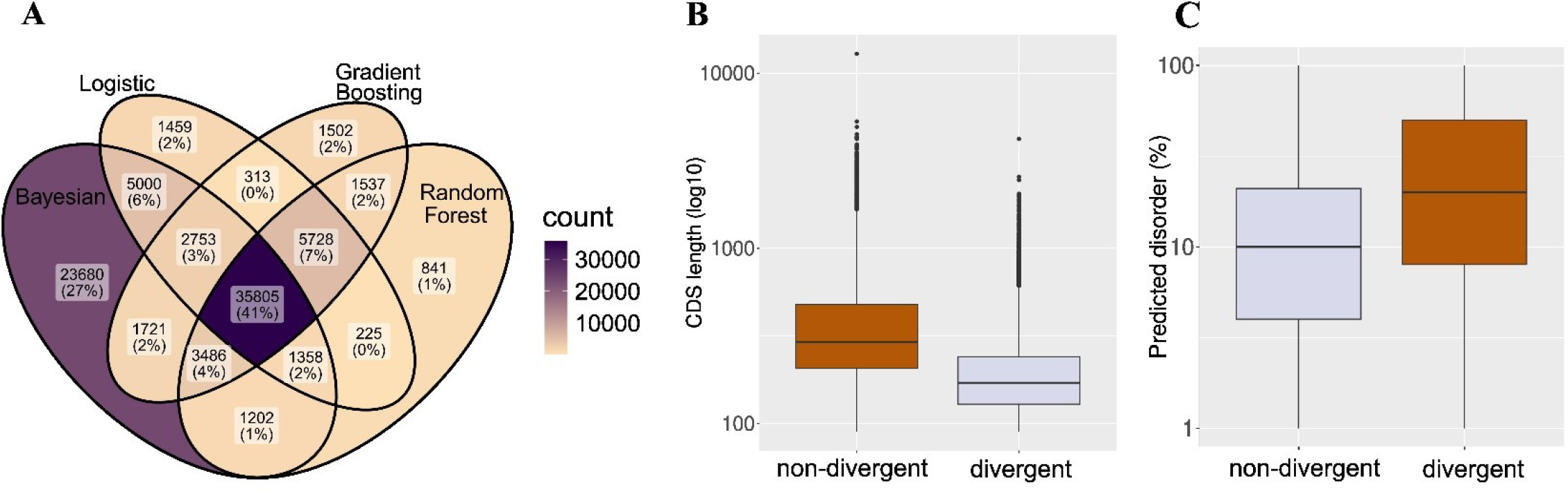
**A**) Overlap of predictions for the divergent (positive) class of the best performing models, on real orphans produced by Vakirlis & Kupczok. **B**) CDS length of predicted non-divergent (n=123,378) and divergent (n=51,205) orphans. **C**) Percentage of intrinsic protein disorder of predicted non-divergent and divergent orphans.

We used each separate simulated orphan dataset together with its negative set of same number of reversed sequences, to train and test four machine learning models using 5-fold cross-validation. Accuracy, as measured on a separate validation set, ranged from 51 to 88.7% (**Figure 2B;** see **Supp. Figure 5** for rest of metrics) with the best performing models being Gradient Boosting and Logistic Regression when trained on sets generated with alpha=0.1, indel rate=0.005 and no invariant sites (accuracy=88.69%; note that compared to the rest of the parameters invariant sites had negligible effect). We found marginally better results when using UHGP100 (54.6-90.95% accuracy; see **Supp. Figure 3B, C**). To confirm that our results are robust to the E-value cut-off of 0.001, we tested an additional set generated with the abovementioned best-performing parameters but defining orphans using an E-value cut-off of 0.01, and we achieved an even higher performance of 97.22% accuracy with Logistic Regression (the rest of the models also scored over 90%). The Bayesian classifier showed consistently poorer performance than the other three. Generally, the best performing models were trained under lower alpha values. Simulating sequences without any insertions or deletions also significantly boosted performance including for sets with higher alpha values (**Figure 2C**; best accuracy reached >95%).

Using only sequences simulated along the two shortest branches of the tree (and thus overall less diverged) to train and test the models improved performance across the board, with the best performing models reaching 91.84% accuracy (**Figure 2D, Supp. Figure 6**). For these best performing models, which we use in downstream analyses, we also controlled for length differences within the training set, since we noticed that simulated orphans were shorter on average than reversed sequences (**Supp. Figure 7**). Indeed a known crucial factor in the efficacy of similarity searches is query sequence length^47^ and short length, rate of divergence and evolutionary age are known to be intertwined as shorter sequences are statistically quicker to fade into the “twilight zone” of sequence similarity^31,47,48^. As this length difference could introduce a classification bias, we truncated the reversed root sequences so that the resulting reversed orphans would have the same average length as simulated ones (see **Methods**). Our findings were overall robust to this adjustment. (**Supp. Figure 8**), and this was also true when using UHGP100 (**Supp. Figure 3D**). Note that features that depend on the length of the query were already normalized by the sequence length (i.e. top bit score, average bit score, alignment count etc.).

As natural sequences are unlikely to evolve under a single parameter scenario, we tested our model on a dataset containing an equal mix of all parameter combinations (using the two shortest branches only and correcting for length differences; see **Methods**), by combining the features of all sets into a single matrix, and achieved an accuracy of 72.24% with Logistic Regression.

In summary, we demonstrate that under sets of evolutionary simulation parameters that favor the persistence of some traces of similarity, simulated diverged orphans can be detected with very high accuracy, while lower but still useful levels of accuracy can be achieved even in more “extreme” divergence scenarios as well as a mix of different scenarios.

As the classic ML models used here are explainable, we next asked which features among the ones we used were the most informative. Feature importance was determined using the permutation approach for the Bayesian, random forest and gradient boosting classification models. When a single feature value is randomly shuffled, the model score drops, and this is what is meant by the permutation feature importance. For the best performing model, the feature that had the most or the second most information consistently in all three models, with the exception of Bayesian, was alignment count **(Figure 2E, Supp. Figure 3E** for UHGP100). Yet, when comparing the distribution of alignment count between the simulated orphans (divergent) and the reversed orphans (non-divergent) we found no significant difference between the two classes (simulated mean=0.025, reversed mean = 0.027; **Figure 2F**). The features that did show the greatest difference were the ones based on identity and E-value (**Figure 2F**). Furthermore, such features showed much lower differences in a model with lower performance (mix of all parameter combinations), suggesting that they are important for correct classification (**Supp. Figure 9**)^49^. Note that alignment count is positively correlated with 5 other features (**Supp. Figure 10**). We conducted additional training and testing excluding alignment count as a feature and the results showed that Logistic Regression maintained performance (89.8% accuracy), whereas Random Forest and Gradient Boosting showed a decrease of 10% in accuracy (77.55% for both). Thus, alignment count, although not significantly different in itself, is nevertheless an impactful feature of the models.

### Thousands of real gut microbiome orphans are predicted to be divergent, and they are shorter and more disordered than non-divergent ones

With a robust classifier at hand, we next moved to test our best performing models (those corresponding to **Figure 2D**) on real orphan sequences. Vakirlis & Kupczok^35^ identified 631,104 species-specific orphan genes out of which 174,583 met the criteria to be used as input to our classifiers (see **Methods**). To increase confidence in our classification, to classify a gene as divergent we required a divergent prediction by least two models (excluding Bayesian due to its poor performance) and this resulted in 51,205 (∼29%) orphans classified as divergent, with overall good overlap among the three methods (**Figure 3A**). In the same article, the authors identified ∼1,000 de novo gene candidates based on conserved synteny and absence of coding potential in the orthologous region of outgroup species. Out of these, 227 were eligible for prediction and 95 (41%) are predicted as diverged. Thus, contrary to what we would expect, we find a significant enrichment in the diverged class among the de novo candidates compared to the remaining orphans (Chi-squared test=16.591, P-value=4.638e-05). This enrichment persists, albeit weaker, when controlling for length differences that exist between de novo candidates and the rest of orphans, by considering only orphans that are no longer than the average length of de novo candidates (Chi-squared test=6.49, P-value=0.0108). This raises the question of whether these genes might have been incorrectly identified as de novo genes in the initial study, or whether they pose a classification challenge for our model which leads to it underperforming in this case.

If our predictions on real orphans reflect their true evolutionary origins, we may expect predicted divergent orphans to differ in some properties compared to non-divergent ones, especially if the non-divergent class is enriched for de novo gene candidates which are known to have some distinguishing properties^23^. We thus compared a number of properties, such as length, GC-content, isoelectric point, intrinsic disorder etc. One property that showed a significant difference was length (**Figure 3B**), which is consistent with our findings about the impact of length on the detection of orphans and on the performance of the models. Additionally, proteins that are predicted to be divergent had a higher intrinsic disorder percentage (**Figure 3C**), consistent with disordered proteins generally evolving faster than globular ones^50^. There were two additional properties for which divergent and non-divergent orphans differed, both minor: the first was the terminal branch length on the bacterial species tree for the specific species a given orphan comes from (shorter for divergent; Cohen’s d=-0.17; Wilcoxon p-value<2.2e-16). The other difference is the *d*_*N*_*/d*_*S*_, with the non-divergent being under slightly weaker negative selection (Cohen’s d = -0.21; Wilcoxon p-value<2.2e-16), which makes sense since non-divergent genes are more likely young de novo genes that are known to be under less constraints^22,26,28,31^. While the rest of the properties showed no statistically significant difference, the ones that do, and especially intrinsic disorder, indicate that our predictions do reflect the divergent origins of these orphans.

## Discussion

Orphan genes constitute a unique and challenging case of evolutionary products. The lack of homologues obscures our insight not only on their origin but also their function, with the exception of well-studied model organisms. The annotation of coding and non-coding genes is a time-consuming, difficult and error-prone process^51–53^ and in combination with the limitations of similarity search algorithms to detect remote homology, it can make characterizing such genes a challenging task.

Here we showed that classic machine learning models can achieve high performance in the task of identifying orphan genes with homologues in the “twilight zone” of sequence similarity, by examining simple statistical features extracted from DIAMOND hits with non-statistically significant E-value. To our knowledge, this is the first attempt at looking for signal within these data which are usually discarded as uninformative. Our results clearly show that ML models have a high discriminative ceiling, but not across the entire range of possible evolutionary scenarios. When at least some sites evolve slower than others and the amount of divergence is not extreme the models perform well, presumably thanks to some residual similarity, in line with what has been observed before but for statistically significant hits^31^. When this isn’t the case, performance drops albeit to still useful levels. The fact that it is hard to assess the mix of evolutionary parameter combinations that underlie the natural evolution of orphan genes is a limitation of our approach which relies on simulations. Yet it is encouraging that even when training is done on a mixed dataset that contains an equal number of orphans from simulations conducted under all possible parameter combinations, the models achieve an accuracy of 72.24%.

Reversed protein sequences serve as a robust null model, yet one potential limitation is that they cannot fully represent de novo originated proteins. Assuming that de novo origination is indeed the main alternative process in real orphan sequence evolution this means that the most appropriate control would be a gold standard set of de novo genes. As this does not exist yet one direction for future work might be to use translated small intergenic ORFs, which can form the basis for de novo gene birth^22,54,55^, as the non-divergent class. This approach would also allow the use of sequence features, including embeddings from protein language models, in the training of more complex machine learning models. Such deep learning techniques have already powered advancements in searches for functional similarity^56^ and the two problems are fundamentally similar. Other worthwhile improvements that future studies could pursue could come from using structural similarity as a comparison and potentially as a benchmark^40,53^. Additionally, versions of this analysis with a wider taxonomic scope (e.g. using the entire UniProt database as target) could leverage the taxonomic distribution of species with similarity hits to test whether a pattern exists in the divergence class, e.g. by giving higher importance to hits coming from species that are phylogenetically closer to the focal species, for which higher similarity might be expected under a divergent scenario.

We believe that the present work shows that simple, computationally inexpensive machine learning can offer valuable insights into evolutionary and comparative genomics problems. Our best performing classifier can be readily and easily used as an additional filter of similarity searches, even if only for sequences that have at least one nonsignificant hit. For example, during analysis of the protein-coding repertoire of a new species, it is not uncommon to find percentages of orphans that exceed 20%^2,24,34^. When that happens, our classifier can provide an immediate estimate of how many of the eligible orphans are the result of divergence beyond recognition thus informing downstream analysis decisions and facilitating the formulation of evolutionary hypotheses.

## Materials & Methods

### Dataset

The sequence dataset used here was obtained from the first version of the Unified Human Gastrointestinal Protein catalog (based on UHGG v1.0) clustered at 50% identity (UHGP50) and at 100% identity (UHGP100)^57^. Species-specific orphan genes previously identified based on the UHGP50 clusters (n=631,104; one representative per cluster), and their nucleotide and amino acid sequences were obtained from ref^35^. The remaining 3,968,227 cluster representatives (non-orphans) were deemed suitable for use as root sequences for simulation. Sequence, structural and evolutionary data for orphans and non-orphans used for Figure 3 and associated analyses were obtained from the supplementary data of ref^35^.

### Sequence evolution simulation

For the sequence evolution simulation, we used the R package phylosim^44^. Out of the available set of non-orphan proteins that we could use as an input, we randomly selected 800,000 proteins with the use of seqtk^58^ (version 1.4-r130-dirty). This subset was split further into 16 sets of 50,000 sequences. This was done in order to use different values for the simulation parameters we used. To define appropriate ranges of values for the different parameters, we conducted some initial test simulation runs and counted the number of orphans they produced. LG was selected as a substitution model. For the shape parameter (alpha value) of the gamma distribution we used the following values: 0.1, 0.3, 0.5 and 1. For the indel rates we tested values 0.005 and 0.01. Lastly, the proportion of invariant sites was set to either 0 (i.e. no invariant sites) or 0.1. As the simulation also required a tree, we chose to use one with 4 terminal branches of lengths 1, 3, 5, 10. We set the inner branches to negligible values. The tree is shown below in newick format.

### (((Taxon1:1, Taxon2:3):0.1,Taxon3:5):0.1,Taxon4:10)

For the simulations without insertions or deletions we used SeqGen^59^, as it was significantly faster than phyloSim. The parameters we used as input to SeqGen were the exact same as the ones in the phyloSim with the exception of invariant sites, as we previously observed that the impact of invariant sites was negligible.

For the positive set of the mixed dataset we used orphans that were simulated on the two shortest branches (Taxon1 and Taxon2) and their negative reversed counterparts were truncated to correct for length differences. We randomly sampled an equal number of simulated and an equal number of reversed orphans from each set. That number was 195 and was defined by the subset which produced the lowest number of simulated orphans, which was that with alpha=0.1, indel = 0.005 and proportion of invariant site = 0.001. In total the mixed set contained 16*195 = 3,120 simulated orphans and 3,120 reversed orphans.

### Orphan identification and Feature extraction

Our negative class dataset was generated by reversing (i.e. reading from end to start) the simulated sequences, that is, the sequences obtained at the end of the simulation. To correct for length differences observed further downstream in the analysis, between the reversed and simulated orphans, in some versions of the analysis we truncated the reversed sequences before they were used as queries for similarity searches, so that the length distributions of the two orphan sequence sets (reversed and simulated) would have similar means before the similarity search is conducted. The percentage of the sequence that was truncated was defined empirically based on the observed differences between reversed and simulated orphans when truncation is not performed.

Both the simulated (positive) and reversed (negative) sequences were aligned to either the UHGP50 or UHGP100 protein sequence dataset with the use of DIAMOND^43^, as shown below:

`diamond blastp -q $path/to/query -d $path/to/database --ultra-sensitive -o $path/to/output -f 6 -p 12 -e #value`

As eligible orphans we characterized the proteins that had no match with E-value≤0.001 and had at least one match with E-value<1. For each eligible orphan, we extracted features from the DIAMOND output to train the classification models. These features are: alignment count (divided by query length), average percent identity, average E-value, average bit score (divided by query length), average coverage (average of alignment length/query length), lowest E-value, highest percent identity, average alignment length, highest bit score (divided by query length), highest alignment length (divided by query length), and total query coverage, and these were calculated by taking into account all the alignments (hits) of each eligible orphan with 0.001 ≤ E-value ≤ 1. Each feature was z-score normalized. In addition, the column “CLASS” was added to denote the origin of the query (target class). The features were then used to train the machine learning models in a supervised manner.

### Classification models

The combined tables were split into training (75%) and test (25%) sets using the Sklearn software^60^. The features in these tables were used to train a bayesian, logistic regression, random forest and gradient boosting classification model from the Sklearn package. For each model, cross-validation was employed to select the optimal parameters. For this research, 5-fold cross-validation was used for all models. For the random forest model, parameters were optimized for the number of estimators (1 to 1000 in steps of 50), max feature function (sqrt or log2) and the criterion function (gini, entropy and log loss). The parameters for the gradient boosting classifier model were optimized for the number of estimators (1, to 1000 in steps of 50), max feature function (sqrt or log2), learning rate (0.1 till 1 in steps of 0.1) and the loss function (log loss and exponential). The Bayesian model was only optimized for the var smoothing parameter for which 1 to 20 was tested in steps of 1. Finally, the built-in cross validation from the Sklearn package was used to optimize the parameters for the logistic regression model.

For each model, accuracy, precision, recall and Matthew’s Correlation Coefficient were calculated. Additionally, the top 10 most important features were selected. The logistic regression model’s features with the greatest and lowest coefficients were chosen, since a large (negative or positive) value indicates that the coefficient has some effect on the prediction. For the Bayesian, random forest and gradient boosting classification models, the feature importance was determined using the permutation approach, which describes whether the model score drops when a single feature value is randomly shuffled.

## Acknowledgements

This work was supported by a PhD scholarship by a Fondation Santé “Sidney Altman Scholarship Program” to Emilios Tassios. The research project was supported by the Hellenic Foundation for Research and Innovation (H.F.R.I.) under the “3rd Call for H.F.R.I. Research Projects to support Post-Doctoral Researchers” to Nikolaos Vakirlis (Project Number:7330) and by a G4 grant from the Pasteur Institute awarded to Nikolaos Vakirlis.

## Data availability

The conda environment with all required modules and packages, the scripts to train the models, and the best pretrained models are available at https://github.com/emiliostassios/Classification-of-divergent-genes-using-ML

