## Supplementary Material for "Machine learning can identify orphans that have diverged into the “twilight zone” of sequence similarity"

A

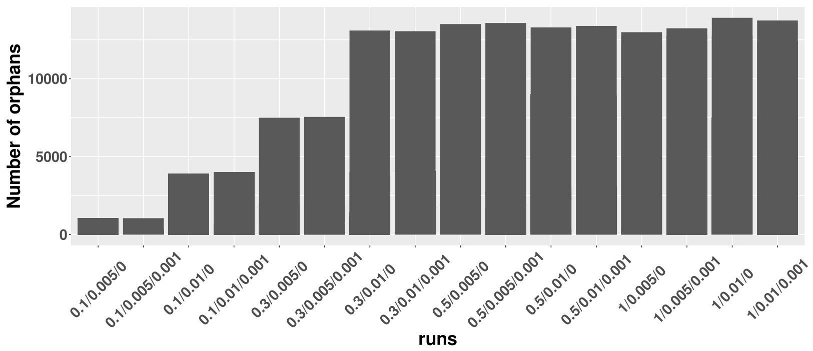


B

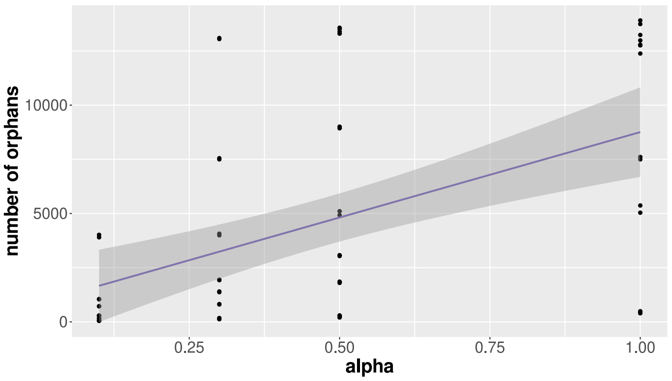


**Supplementary Figure 1**: **A**) The number of simulated orphans per parameter set. Parameter values shown on x-axis: alpha/indel rate/proportion of invariant sites. **B**) Correlation of the number of simulated orphans to the alpha value used in the simulation (Spearman's Rho = 0.55; P-value = 2e-06).

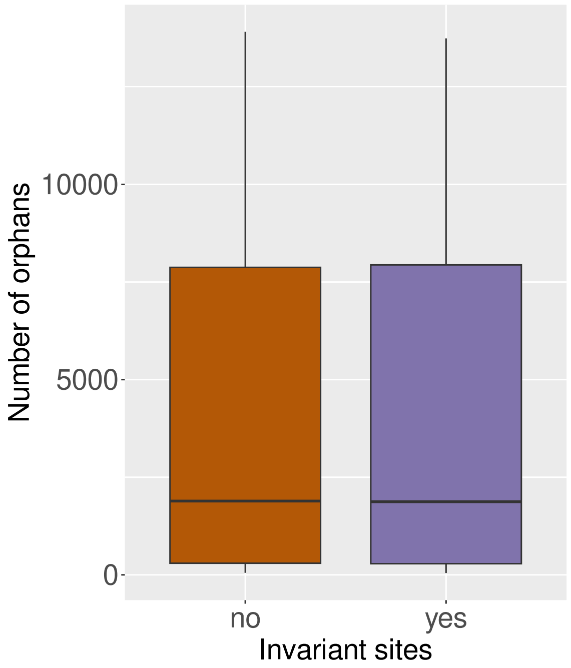


**Supplementary Figure 2:** Number of simulated orphans for sets that were simulated with and without invariant sites. The parameter seems to not be important in the number of simulated orphans produced.

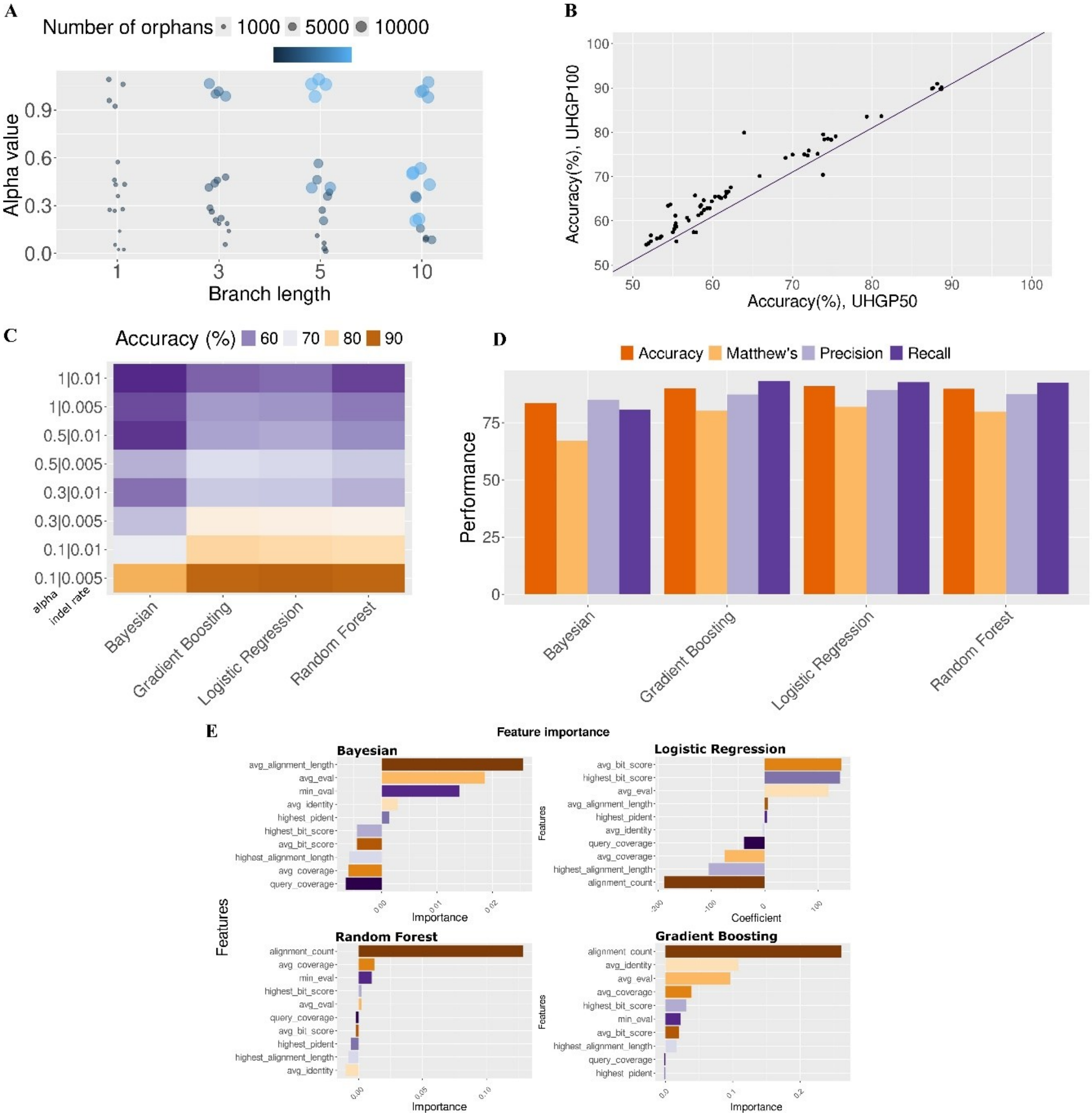


**Supplementary Figure 3**: Equivalent to Figure 2 but using UHGP100 as a target for similarity searches. **A**) Scatterplot showing the correlation of accuracies across the different combinations of simulation parameters, when using the UHGP50 (x-axis) or UHGP100 (y-axis) for similarity searches. Across the range of parameters, searching within the larger database produces marginally better performance. **B**) Number of simulated orphans per alpha value and branch length. **C**) Accuracy metric for all four models (Bayesian, Logistic regression, Gradient Boosting and Random Forest) for all subsets produced with various combinations of alpha and indel rate values, without invariant sites. **D**) All performance metrics (accuracy, recall, precision, Matthew's correlation coefficient) for the best performing models, as shown in C (alpha=0.1, indel rate=0.005). **E**) The most important features for each one of the best performing models.

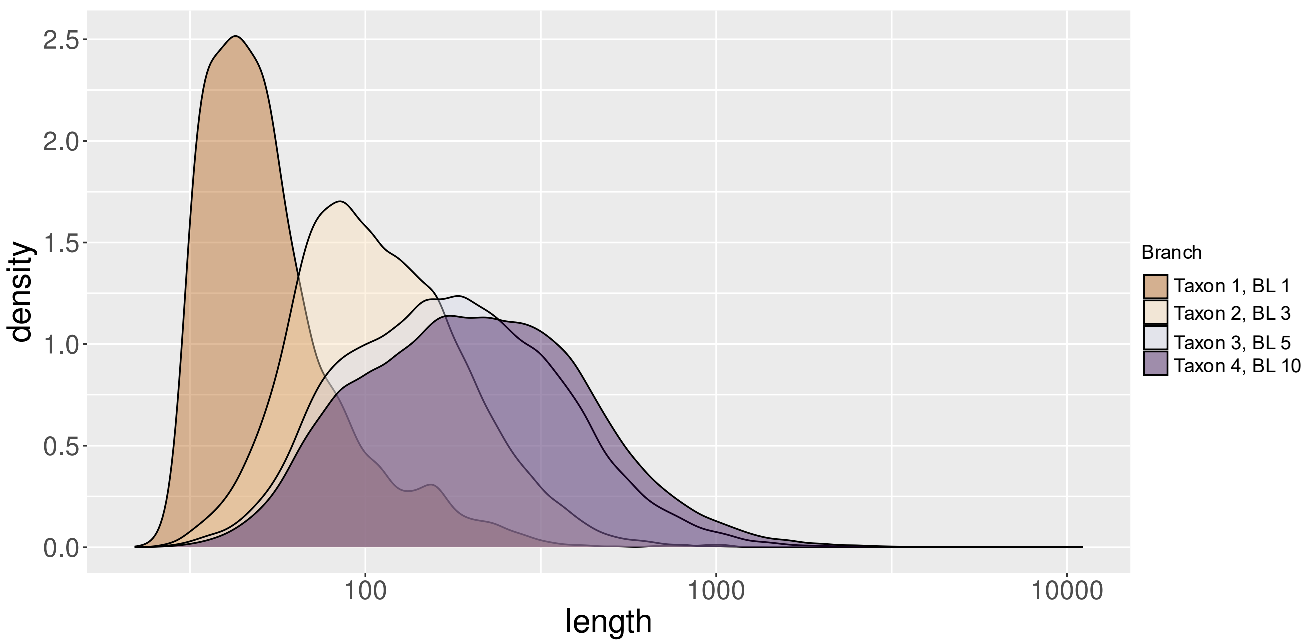


**Supplementary Figure 4**: The length distribution of the simulated orphans produced on different branches. The orphans simulated on the shortest branches were shorter.

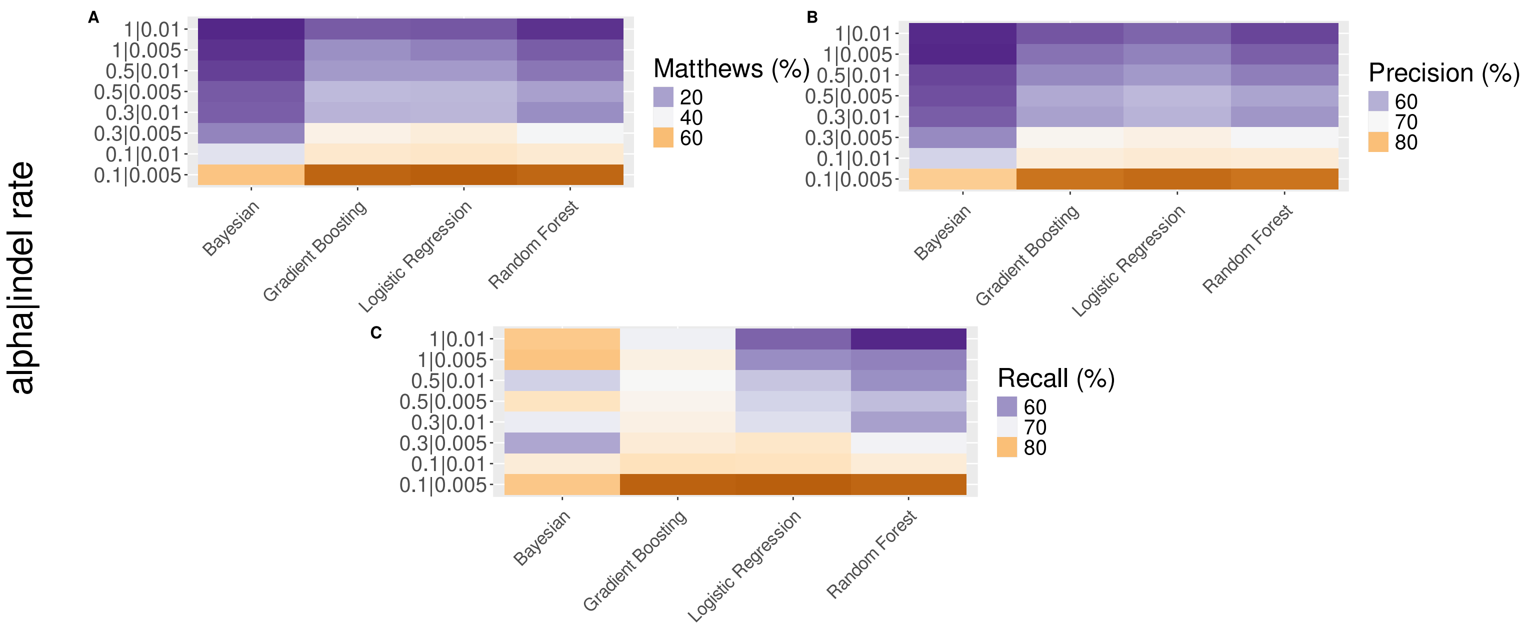


**Supplementary Figure 5**: The rest of the metrics for the performance of the models for combinations of alpha and indel rate parameters. **A**) Matthew's correlation coefficient, **B**) precision, **C**) recall.

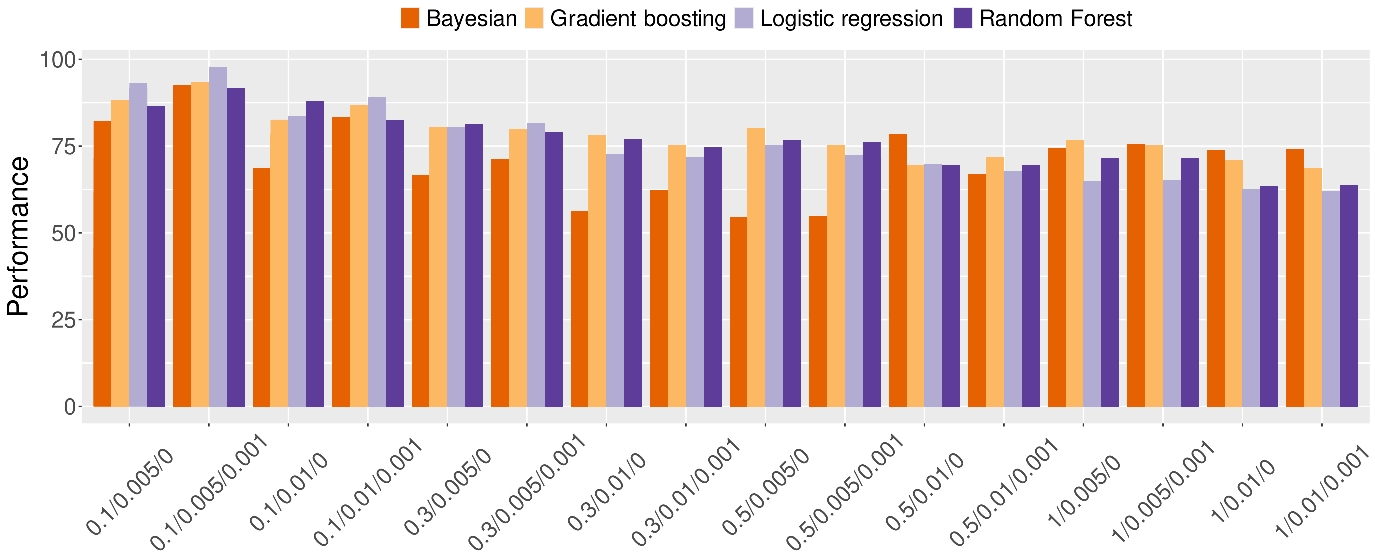


**Supplementary Figure 6**: Accuracy (as a percentage) for all models and parameter combinations, shown on the x-axis as alpha/indel/inv_sites, correcting for length differences between reversed negative controls and simulated orphans and using only sequences simulated along the two shorter branches of the tree.

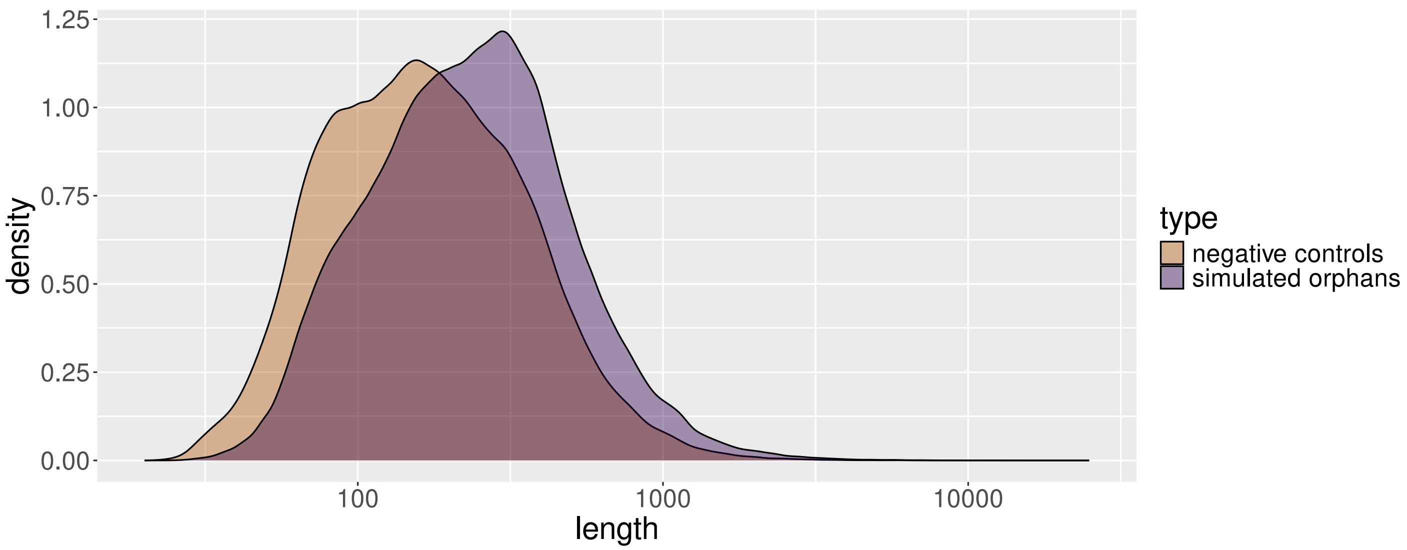


**Supplementary Figure 7**: The length distribution of the simulated orphans and the reversed sequences, that were used as negative controls. The simulated orphans are slightly shorter than their reversed counterparts.

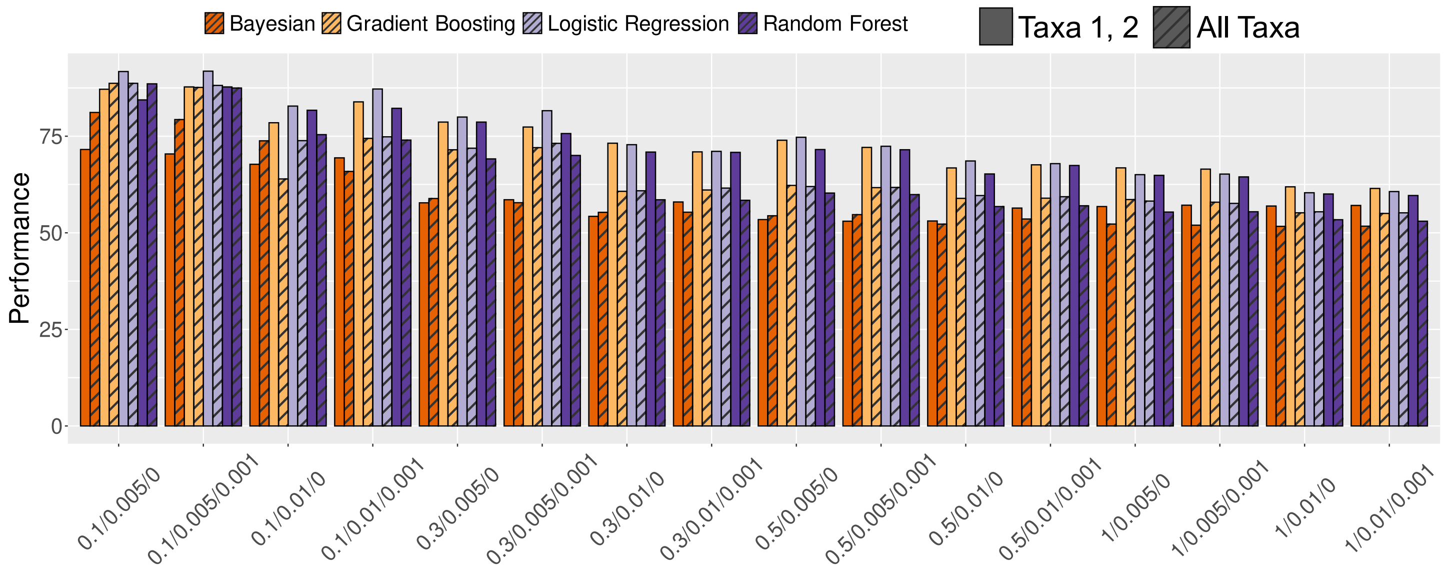


**Supplementary Figure 8**: Accuracy (as a percentage) for all models and parameter combinations, shown on the x-axis as alpha/indel/inv_sites, comparing two versions: solid bars show version where we correct for length differences between reversed negative controls and simulated orphans and use only sequences simulated along the two shorter branches of the tree and striped bars show version using sequences simulated on all four branches and without length correction.

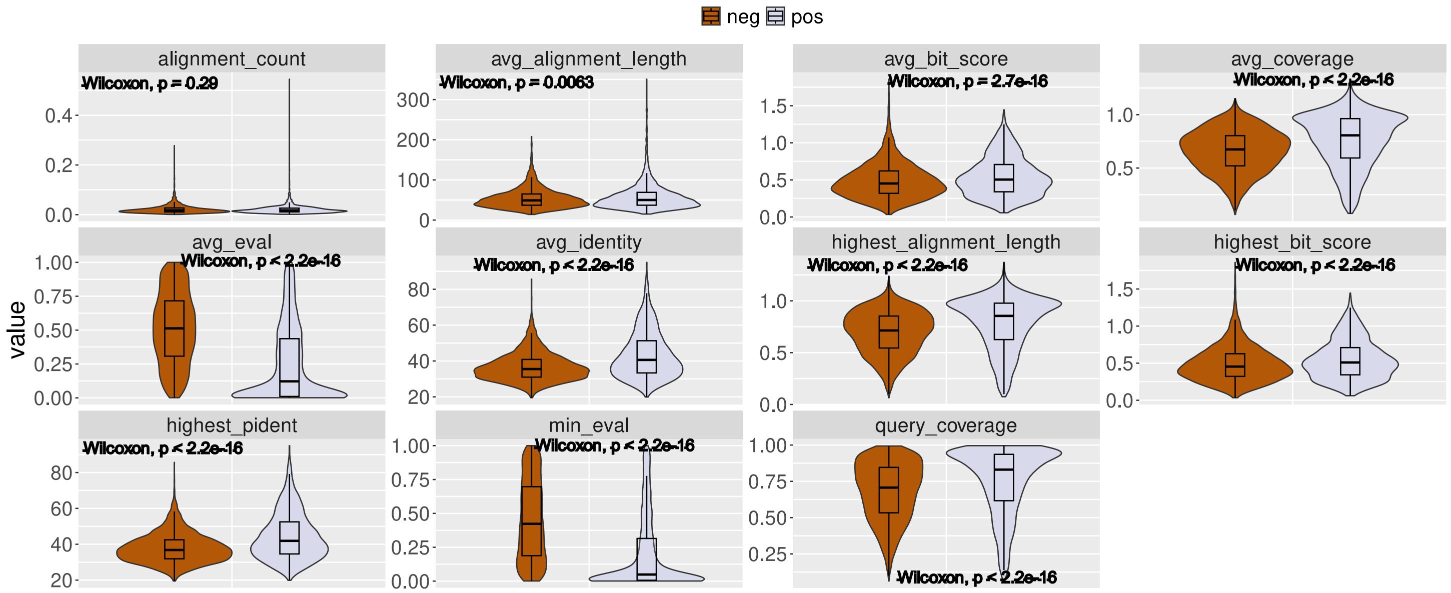


**Supplementary Figure 9***:* The comparison between simulated orphans and reversed sequences of the means of the features used for the training of the mixed set that contained a balanced number of all the different subsets simulated under different parameter combinations.

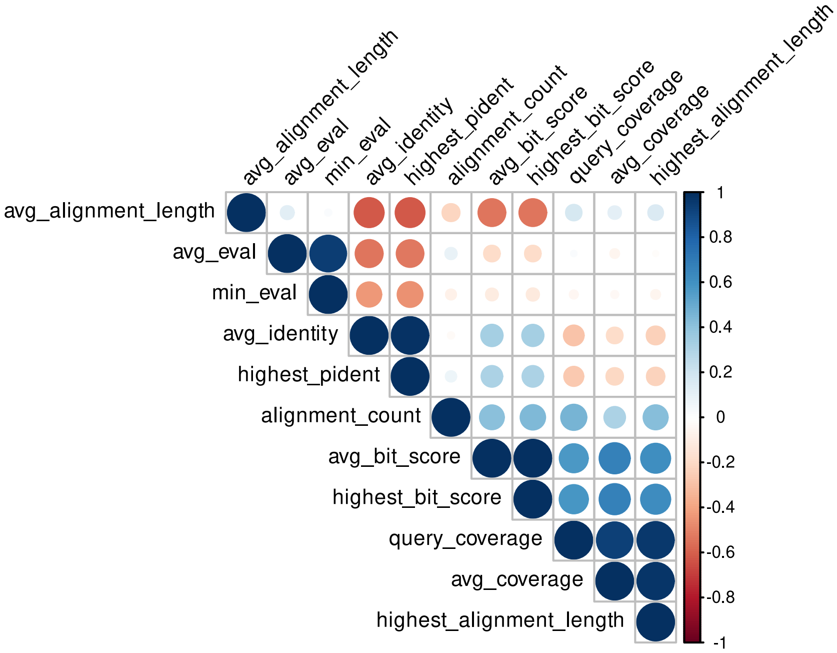


**Supplementary Figure 10:** Correlation matrix of the features used to train the best performing model (Logistic Regression trained with simulation parameters alpha = 0.1, indel = 0.005, inv_sites = 0.001; only simulated orphans from the two shortest branches used), after correcting for length differences between simulated and reversed orphans.

**Supplementary Table 1**: The performance measured by accuracy of all the models with all of the different parameter combinations.

| **model** | **alpha** | **indel_rate** | **invariant_sites** | **accuracy** |
|---|---|---|---|---|
| Bayesian | 0.1 | 0.005 | 0 | 81.14 |
| Gradient Boosting | 0.1 | 0.005 | 0 | 88.69 |
| Random Forest | 0.1 | 0.005 | 0 | 88.56 |
| Logistic Regression | 0.1 | 0.005 | 0 | 88.69 |
| Bayesian | 0.1 | 0.005 | 0.001 | 79.31 |
| Gradient Boosting | 0.1 | 0.005 | 0.001 | 87.62 |
| Random Forest | 0.1 | 0.005 | 0.001 | 87.48 |
| Logistic Regression | 0.1 | 0.005 | 0.001 | 88.14 |
| Bayesian | 0.1 | 0.01 | 0 | 73.81 |
| Gradient Boosting | 0.1 | 0.01 | 0 | 63.94 |
| Random Forest | 0.1 | 0.01 | 0 | 75.41 |
| Logistic Regression | 0.1 | 0.01 | 0 | 73.85 |
| Bayesian | 0.1 | 0.01 | 0.001 | 65.88 |
| Gradient Boosting | 0.1 | 0.01 | 0.001 | 74.43 |
| Random Forest | 0.1 | 0.01 | 0.001 | 74 |
| Logistic Regression | 0.1 | 0.01 | 0.001 | 74.85 |
| Bayesian | 0.3 | 0.005 | 0 | 58.87 |
| Gradient Boosting | 0.3 | 0.005 | 0 | 71.49 |
| Random Forest | 0.3 | 0.005 | 0 | 69.13 |
| Logistic Regression | 0.3 | 0.005 | 0 | 71.9 |
| Bayesian | 0.3 | 0.005 | 0.001 | 57.79 |
| Gradient Boosting | 0.3 | 0.005 | 0.001 | 72.04 |
| Random Forest | 0.3 | 0.005 | 0.001 | 70.02 |
| Logistic Regression | 0.3 | 0.005 | 0.001 | 73.15 |
| Bayesian | 0.3 | 0.01 | 0 | 55.3 |
| Gradient Boosting | 0.3 | 0.01 | 0 | 60.74 |
| Random Forest | 0.3 | 0.01 | 0 | 58.55 |

| | | | | |
|---|---|---|---|---|
| Logistic Regression | 0.3 | 0.01 | 0 | 60.88 |
| Bayesian | 0.3 | 0.01 | 0.001 | 55.33 |
| Gradient Boosting | 0.3 | 0.01 | 0.001 | 61.08 |
| Random Forest | 0.3 | 0.01 | 0.001 | 58.42 |
| Logistic Regression | 0.3 | 0.01 | 0.001 | 61.57 |
| Bayesian | 0.5 | 0.005 | 0 | 54.41 |
| Gradient Boosting | 0.5 | 0.005 | 0 | 62.27 |
| Random Forest | 0.5 | 0.005 | 0 | 60.29 |
| Logistic Regression | 0.5 | 0.005 | 0 | 61.97 |
| Bayesian | 0.5 | 0.005 | 0.001 | 54.7 |
| Gradient Boosting | 0.5 | 0.005 | 0.001 | 61.69 |
| Random Forest | 0.5 | 0.005 | 0.001 | 59.89 |
| Logistic Regression | 0.5 | 0.005 | 0.001 | 61.74 |
| Bayesian | 0.5 | 0.01 | 0 | 52.25 |
| Gradient Boosting | 0.5 | 0.01 | 0 | 58.91 |
| Random Forest | 0.5 | 0.01 | 0 | 56.79 |
| Logistic Regression | 0.5 | 0.01 | 0 | 59.64 |
| Bayesian | 0.5 | 0.01 | 0.001 | 53.57 |
| Gradient Boosting | 0.5 | 0.01 | 0.001 | 58.96 |
| Random Forest | 0.5 | 0.01 | 0.001 | 57 |
| Logistic Regression | 0.5 | 0.01 | 0.001 | 59.34 |
| Bayesian | 1 | 0.005 | 0 | 52.26 |
| Gradient Boosting | 1 | 0.005 | 0 | 58.6 |
| Random Forest | 1 | 0.005 | 0 | 55.37 |
| Logistic Regression | 1 | 0.005 | 0 | 58.21 |
| Bayesian | 1 | 0.005 | 0.001 | 51.97 |
| Gradient Boosting | 1 | 0.005 | 0.001 | 57.92 |
| Random Forest | 1 | 0.005 | 0.001 | 55.45 |
| Logistic Regression | 1 | 0.005 | 0.001 | 57.61 |
| Bayesian | 1 | 0.01 | 0 | 51.67 |
| Gradient Boosting | 1 | 0.01 | 0 | 55.18 |
| Random Forest | 1 | 0.01 | 0 | 53.39 |
| Logistic Regression | 1 | 0.01 | 0 | 55.44 |
| Bayesian | 1 | 0.01 | 0.001 | 51.71 |

| | | | | |
|---|---|---|---|---|
| Gradient Boosting | 1 | 0.01 | 0.001 | 55.01 |
| Random Forest | 1 | 0.01 | 0.001 | 53 |
| Logistic Regression | 1 | 0.01 | 0.001 | 55.17 |

**Supplementary Table 2***:* The performance measured by accuracy of all the models for the orphans that were simulated on the two shortest branches and the reversed sequences, used as negative controls, were length corrected.

| **model** | **alpha** | **indel_rate** | **invariant_sites** | **accuracy** |
|---|---|---|---|---|
| Bayesian | 0.1 | 0.01 | 0.001 | 69.4 |
| Gradient boosting | 0.1 | 0.01 | 0.001 | 83.89 |
| Logistic regression | 0.1 | 0.01 | 0.001 | 87.22 |
| Random Forest | 0.1 | 0.01 | 0.001 | 82.22 |
| Bayesian | 0.1 | 0.01 | 0 | 67.74 |
| Gradient boosting | 0.1 | 0.01 | 0 | 78.49 |
| Logistic regression | 0.1 | 0.01 | 0 | 82.8 |
| Random Forest | 0.1 | 0.01 | 0 | 81.72 |
| Bayesian | 0.1 | 0.005 | 0.001 | 70.41 |

| | | | | |
|---|---|---|---|---|
| Gradient boosting | 0.1 | 0.005 | 0.001 | 87.76 |
| Logistic regression | 0.1 | 0.005 | 0.001 | 91.84 |
| Random Forest | 0.1 | 0.005 | 0.001 | 87.76 |
| Bayesian | 0.1 | 0.005 | 0 | 71.56 |
| Gradient boosting | 0.1 | 0.005 | 0 | 87.16 |
| Logistic regression | 0.1 | 0.005 | 0 | 91.74 |
| Random Forest | 0.1 | 0.005 | 0 | 84.4 |
| Bayesian | 0.3 | 0.01 | 0.001 | 57.98 |
| Gradient boosting | 0.3 | 0.01 | 0.001 | 70.95 |
| Logistic regression | 0.3 | 0.01 | 0.001 | 71.08 |
| Random Forest | 0.3 | 0.01 | 0.001 | 70.82 |
| Bayesian | 0.3 | 0.01 | 0 | 54.26 |
| Gradient boosting | 0.3 | 0.01 | 0 | 73.19 |
| Logistic regression | 0.3 | 0.01 | 0 | 72.81 |
| Random Forest | 0.3 | 0.01 | 0 | 70.9 |
| Bayesian | 0.3 | 0.005 | 0.001 | 58.56 |
| Gradient boosting | 0.3 | 0.005 | 0.001 | 77.38 |
| Logistic regression | 0.3 | 0.005 | 0.001 | 81.61 |
| Random Forest | 0.3 | 0.005 | 0.001 | 75.69 |
| Bayesian | 0.3 | 0.005 | 0 | 57.76 |
| Gradient boosting | 0.3 | 0.005 | 0 | 78.66 |
| Logistic regression | 0.3 | 0.005 | 0 | 79.96 |
| Random Forest | 0.3 | 0.005 | 0 | 78.66 |
| Bayesian | 0.5 | 0.01 | 0.001 | 56.4 |
| Gradient boosting | 0.5 | 0.01 | 0.001 | 67.6 |
| Logistic regression | 0.5 | 0.01 | 0.001 | 67.9 |
| Random Forest | 0.5 | 0.01 | 0.001 | 67.42 |
| Bayesian | 0.5 | 0.01 | 0 | 53.06 |
| Gradient boosting | 0.5 | 0.01 | 0 | 66.79 |
| Logistic regression | 0.5 | 0.01 | 0 | 68.59 |
| Random Forest | 0.5 | 0.01 | 0 | 65.23 |
| Bayesian | 0.5 | 0.005 | 0.001 | 53 |
| Gradient boosting | 0.5 | 0.005 | 0.001 | 72.1 |
| Logistic regression | 0.5 | 0.005 | 0.001 | 72.4 |

| | | | | |
|---|---|---|---|---|
| Random Forest | 0.5 | 0.005 | 0.001 | 71.5 |
| Bayesian | 0.5 | 0.005 | 0 | 53.42 |
| Gradient boosting | 0.5 | 0.005 | 0 | 73.96 |
| Logistic regression | 0.5 | 0.005 | 0 | 74.73 |
| Random Forest | 0.5 | 0.005 | 0 | 71.55 |
| Bayesian | 1 | 0.01 | 0.001 | 57.08 |
| Gradient boosting | 1 | 0.01 | 0.001 | 61.48 |
| Logistic regression | 1 | 0.01 | 0.001 | 60.67 |
| Random Forest | 1 | 0.01 | 0.001 | 59.65 |
| Bayesian | 1 | 0.01 | 0 | 56.93 |
| Gradient boosting | 1 | 0.01 | 0 | 61.9 |
| Logistic regression | 1 | 0.01 | 0 | 60.37 |
| Random Forest | 1 | 0.01 | 0 | 60.05 |
| Bayesian | 1 | 0.005 | 0.001 | 57.15 |
| Gradient boosting | 1 | 0.005 | 0.001 | 66.48 |
| Logistic regression | 1 | 0.005 | 0.001 | 65.2 |
| Random Forest | 1 | 0.005 | 0.001 | 64.47 |
| Bayesian | 1 | 0.005 | 0 | 56.79 |
| Gradient boosting | 1 | 0.005 | 0 | 66.81 |
| Logistic regression | 1 | 0.005 | 0 | 65.05 |
| Random Forest | 1 | 0.005 | 0 | 64.87 |
